# YieldNet: A Convolutional Neural Network for Simultaneous Corn and Soybean Yield Prediction Based on Remote Sensing Data

**DOI:** 10.1101/2020.12.05.413203

**Authors:** Saeed Khaki, Hieu Pham, Lizhi Wang

**Affiliations:** Industrial Engineering Department, Iowa State University, Ames, Iowa 50011, USA; Syngenta, Slater, Iowa 50244, USA

**Keywords:** Crop yield, Remote Sensing, Deep learning, Transfer learning

## Abstract

Large scale crop yield estimation is, in part, made possible due to the availability of remote sensing data allowing for the continuous monitoring of crops throughout its growth state. Having this information allows stakeholders the ability to make real-time decisions to maximize yield potential. Although various models exist that predict yield from remote sensing data, there currently does not exist an approach that can estimate yield for multiple crops simultaneously, and thus leads to more accurate predictions. A model that predicts yield of multiple crops and concurrently considers the interaction between multiple crop’s yield. We propose a new model called YieldNet which utilizes a novel deep learning framework that uses transfer learning between corn and soybean yield predictions by sharing the weights of the backbone feature extractor. Additionally, to consider the multi-target response variable, we propose a new loss function. Numerical results demonstrate that our proposed method accurately predicts yield from one to four months before the harvest, and is competitive to other state-of-the-art approaches.

## 1 Introduction

The use of satellites and other remote sensing mechanisms has proven to be vital in monitoring the growth of various crops both spatially and temporally [1]. Common uses of remote sensing are to extract various descriptive indices such as normalized difference vegetation index (NDVI), temperature condition index, enhanced vegetation index, and leaf area index [2]. This information can be used for drought monitoring, detecting excessive soil wetness, quantifying weather impacts on vegetation, the evaluation of vegetation health and productivity, and for crop yield forecasting [3, 4, 5]. Moreover, large scale collection of environmental descriptors such as surface temperature and precipitation can also be identified from satellite images [6, 7].

Accurate and non-destructive identification of crop yield throughout a crop’s growth stage enables farmers, commercial breeding organizations, and government agencies to make decisions to maximize yield output, ultimately for a nation’s economic benefit. Early estimations of yield at the field/farm level plays a vital role in crop management in terms of site specific decisions to maximize a crop’s potential. Numerous approaches have been applied for crop yield prediction, such as manual surveys, agro-meteorological models, and remote sensing based methods [8]. However, given the size of a farming operations, manual field surveys are neither efficient nor practical. Moreover, modeling approaches have limitations due to the difficulty in determining model parameters, the scalability to large areas, and the computational resources neccessary [9].

Recently, satellite data has become widely available in various spatial, temporal and spectral resolutions and can be used to predict crop yield over different geographical locations and scales [10]. The availability of Earth observation (EO) data has created new ways for efficient, large-scale agricultural mapping [11]. EO enables a unique mechanism to capture crop information over large areas with regular updates to create maps of crop production and yield. Nevertheless, due to the high spatial resolution needed for accurate yield predictions, Unmanned Aerial Vehicles (UAV) have been promoted for data acquisition [12]. Although UAV platforms have demonstrated superior image capturing abilities, without the assistance of a large workforce, accurately measuring yield for large regions is not feasible.

Remote sensing capabilities have been extensively used for estimating crop yield around the world in various scales (field, county, state). Various studies have been performed using various vegetation indices to estimate yield in maize, wheat, grapes, rice, corn, and soybeans using random forests, neural networks, multiple linear regression, partial least squares regression, and crop models [13, 14, 15, 16, 17, 18, 19]. These research papers showcase the potential, power, and accuracy remote sensing has on estimating yield at a large scale.

For the purpose of this paper, we consider two main crops in the Midwest United States (U.S.), corn (Zea *may*s L.) and soybeans (*Glycine max*). According to the United States Department of Agriculture, in 2019, 89.7 million and 76.1 million acres of corn and soybean were planted, respectively. The combination of these two crops make up approximately 20% of active U.S. farmland [20]. Given the shear size of these farming operations combined with the growing population, actions must be taken to maximize yield. With the help of remote sensing, government officials as well as farmers can monitor crop growth (at various scales) to ensure proper crop management decisions are made to maximize yield potential.

Although approaches exist to estimate yield with various means, current models are limited in that they only estimate yield for a single crop. That is, there does not exist any model making use of remote sensing data to predict the yield of multiple crops simultaneously. Having such a model would result in more accurate predictions by considering the interactions between crops and estimates yields accordingly. Pragmatically, because predictions are simultaneous, the results are less computationally intensive than computing a model for each crop.

We propose a new deep learning framework called YieldNet utilizing a novel deep neural network architecture. This model makes use of a transfer learning methodology that acts as a bridge to share model weights for the backbone feature extractor. Moreover, because of the uniqueness associated to simultaneous yield prediction for two crops, we propose a new loss function that can handle multiple response variables.

## 2 Methodology

The goal of this paper is to simultaneously predict the average yield per unit area of two different crops (corn and soybean) both grown in regions of interest (U.S. counties) based on a sequence of images taken by satellite before harvest. Let 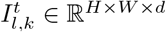 and 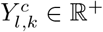 denote, respectively, the remotely sensed multi-spectral image taken at time *t* ∈ {1, …, *T*} during the growing season and the ground truth average yield of crop type *c* ∈ {1, 2} planted in location *l* at year *k*, where *H* and *W* are the image’s height and width and *d* is the number of bands (channels). Thus, the dataset can be represented as

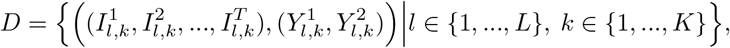

where *L* and *K* are the numbers of locations and years, respectively.

For our approach, we do not use end-to-end training due to the main following reasons: (1) the number of labeled training data is limited, and (2) inability to use transfer learning from popular benchmark datasets such as Imagenet due to domain difference and multi-spectral nature of satellite images. Therefore, we reduce the dimensionality of the raw remote sensing images under permutation invariance assumption which states the average yield mostly depends on the number of different pixel types rather than the position of the pixels in images due to infeasibility of end-to-end training. Similar approaches have been used in other studies [21, 22]. As a result, we separately discretize the pixel values for each band of a multi-spectral image 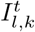 into *b* bins and obtain a histogram representation 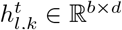. If we obtain histogram representations of the sequence of multi-spectral images denoted as 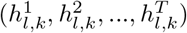 and concatenate them through time dimension, we can produce a compact histogram representations *H*_*l,k*_ ∈ ℝ^*T* ×*b*×*d*^ of the sequence of multi-spectral images taken during growing season. As such, the dataset D can be re-written as the following and we will use this notation in the rest of the paper:

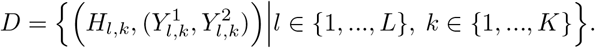

Given the above dataset, this paper proposes a deep learning based method, named YieldNet, that learns the desired mapping 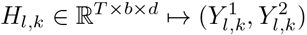 to predict the yields of two different crops simultaneously.

### 2.1 Network Architecture

Crop yield is a highly complex trait which is affected by many factors such as environmental conditions and crop genotype which requires a complex model to reveal the functional relationship between these interactive factors and crop yield.

As such, we propose a convolutional neural network model which is a highly non-linear and complex model. Convolutional neural networks belong to the class of representation learning methods which automatically extract necessary features from raw data without the need of any handcrafted features [23, 24]. Figure 1 outlines the architecture of the proposed method. Table 1 shows the detailed architecture of the proposed model. We use 2-D convolution operation which is performed over the ‘time’ and ‘bin’ dimensions while considering bands as channels. As such, convolution operation over the time dimension can help capture the temporal effect of satellite images collected over time intervals.

**Table 1:** YieldNet architecture. The first 5 convolutional layers work as a backbone feature extractor and share weights for both corn and soybean yield predictions.

| Type/Stride | Padding | Filter Size | Number of Filters |
| --- | --- | --- | --- |
| Conv/s2 | valid | $7 \times 7$ | 48 |
| Conv/s2 | valid | $5 \times 5$ | 64 |
| Conv/s2 | same | $5 \times 5$ | 96 |
| Conv/s1 | same | $3 \times 3$ | 128 |
| Conv/s1 | same | $3 \times 3$ | 128 |
| Corn Head |  | Soybean Head |  |
| Conv/s1 - same - $3 \times 3$ - 148 | | Conv/s1 - same - $3 \times 3$ - 148 | |
| Conv/s1 - same - $3 \times 3$ - 148 | | Conv/s1 - same - $3 \times 3$ - 148 | |
| FC-100 |  | FC-100 |  |
| FC-50 |  | FC-50 |  |

**Figure 1.**
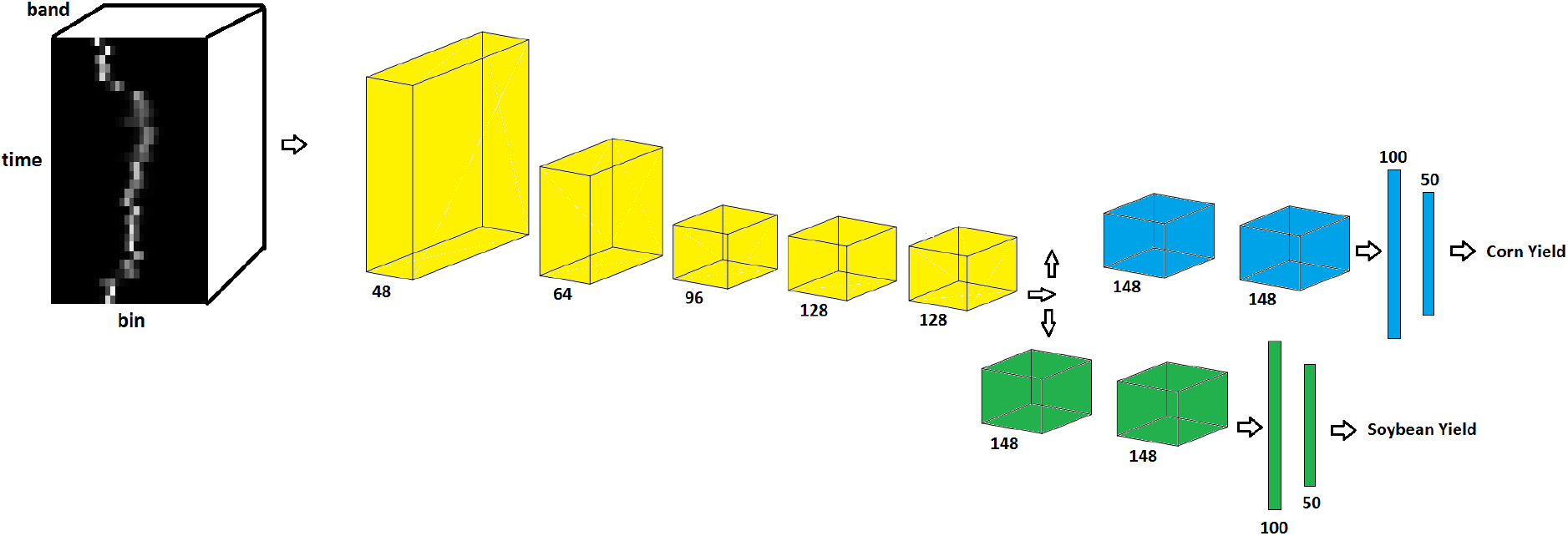
Outline of the YieldNet architecture. The input of the YieldNet is 3-D histograms *H* ∈ ℝ^*T* ×*b*×*d*^ and the 2-D convolution operation is performed over the ‘time’ and ‘bin’ dimensions while considering bands as channels. The number under feature maps indicate the number of channels. Convolutional layers denoted with yellow color work as a backbone feature extractor and share weights for both corn and soybean yield predictions.

In our proposed network, the first 5 convolutional layers share weights for corn and soybean yield predictions which are denoted with yellow color in Figure 1. These layers extract relevant features from input data for both corn and soybean yield predictions. The intuition behind using one common feature extractor is that it significantly decreases the number of parameters of the network, which helps training the model more efficiently given the scarcity of the labeled data. In addition, many low-level features captured by the comment feature extractor reflect general enviromental conditions that are transferable between corn and soybean yields. All convolutional layers are followed by batch normalization [25] and ReLU nonlinearities in our proposed network. Batch normalization is used to accelerate the training process by reducing internal covariate shift and regularizing the network. We use two convolutional layers in both corn and soybean heads which are followed by two fully connected layers.

### 2.2 Network Loss

To jointly minimize the prediction errors of corn and soybean yield forecasting, we propose the following loss function:

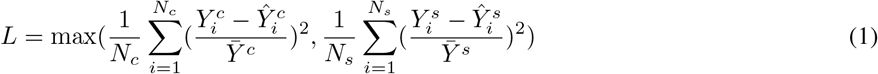

Where 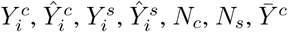 and 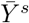 denote *i*th average ground truth corn yield, *i*th predicted corn yield, *i*th average ground truth soybean yield, *i*th predicted soybean yield, number of corn samples, number of soybean samples, average corn yield, and average soybean yield, respectively. Our proposed loss function is a normalized Euclidean loss which makes the corn and soybean losses the same scale. We use the maximum function in our proposed loss function to make the training process more stable and ensure that both corn and soybean losses are optimized.

## 3 Experiments and Results

In this section, we present the dataset used in our study and then report the results of our proposed method along with other competing methods in corn and soybean yield predictions. We conducted all experiments in Tensorflow [26] on a NVIDIA Tesla V100 GPU.

### 3.1 DATA

The data analyzed in this study included two sets: yield performance and satellite images.

- The yield performance dataset includes the observed county-level average yield for corn and soybean between 2004 and 2018 across 1,132 counties for corn and 1,076 for soybean within 13 states of the U.S. Corn Belt: Indiana, Illinois, Iowa, Minnesota, Missouri, Nebraska, Kansas, North Dakota, South Dakota, Ohio, Kentucky, Michigan, and Wisconsin, where corn and soybean are considered the dominant crops [27]. Figures 2 and 4 depict the US Corn Belt and histogram of yields, respectively. The summary statistics of the corn and soybean yields are shown in Table 2.
- Satellite data contains MODIS products including MOD09A1 and MYD11A2. MOD09A1 product provides an estimate of the surface spectral reflectance of Terra MODIS bands 1-7 at 500m resolution and corrected for atmospheric conditions [28]. The MYD11A2 product provides an average land surface temperature which has the day and night-time surface temperature bands at 1km resolution [29]. Figure 3 depicts an example multispectral image of land surface temperature and surface spectral reflectance for Adams county in Illinois. These satellite images were captured at 8-days intervals and we only use satellite images captured during growing seasons (March to October). As such, satellite images are collected 30 times a year in our study. We discretize all multispectral images using 32 bins to generate the 3-D histograms *H* ∈ ℝ ^*T* ×*b*×*d*^, where *T* = 30, *b* = 32, and *d* = 9.
- USDA-NASS cropland data layers (CDL) is a crop-specific land cover data which are produced annually for different crops based on moderate resolution satellite imagery and extensive agricultural ground truth [30]. In this paper, cropland data layers are used for both corn and soybean to focus on only croplands within each county and exclude non-croplands such as buildings and streets from satellite images.

**Table 2:**
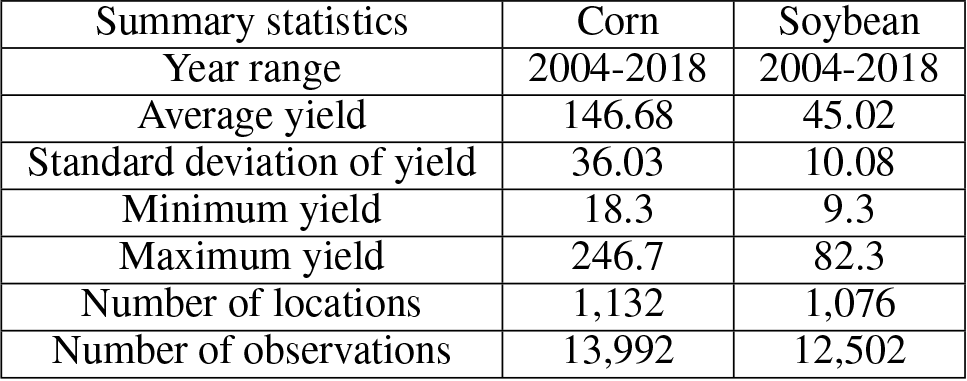
The summary statistics of yield data across all years. The unit of yield is bushels per acre.

**Figure 2.**
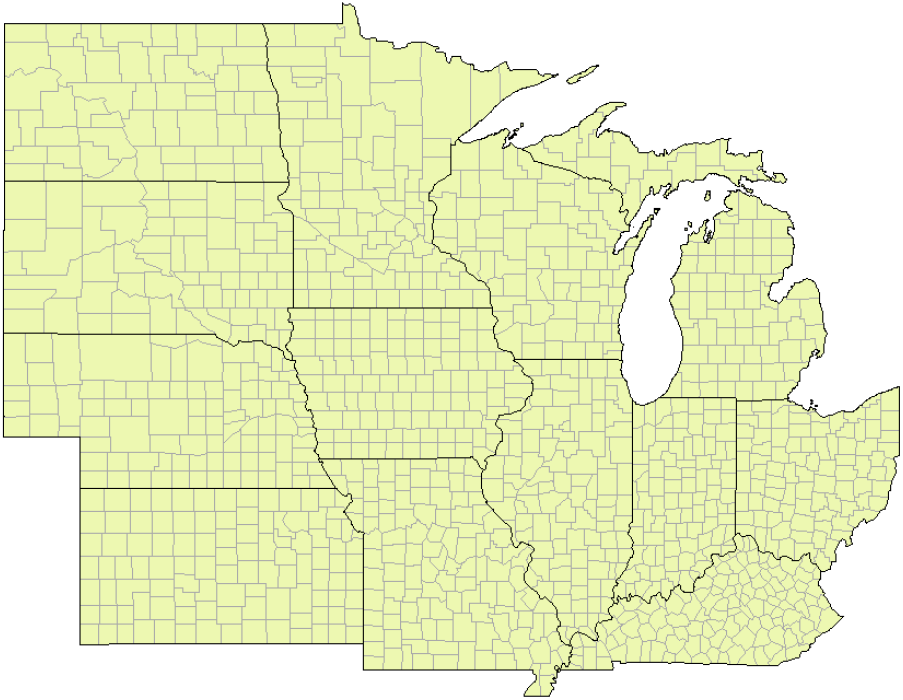
The map of US Corn Belt.

**Figure 3.**
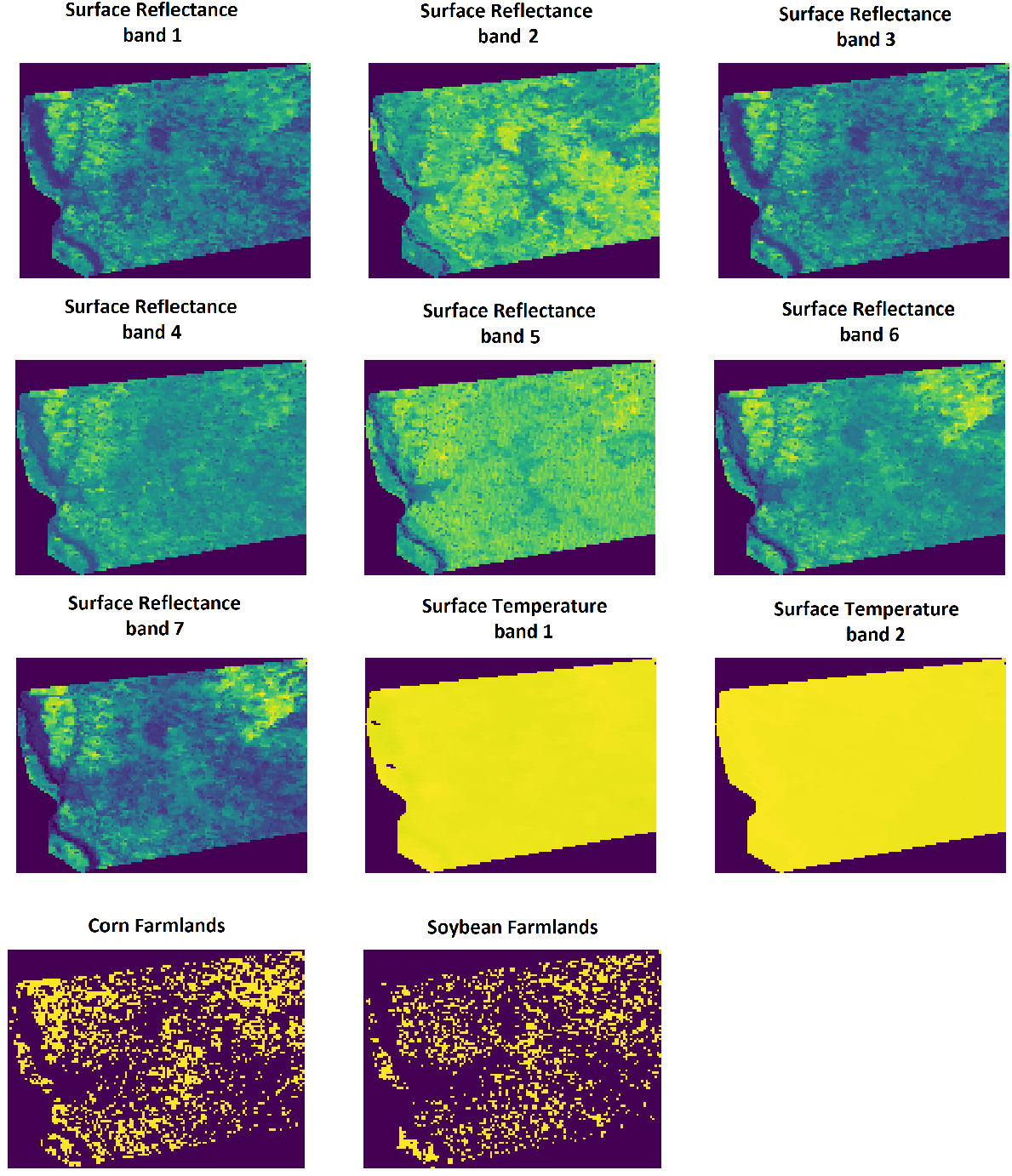
The multispectral images of surface spectral reflectance and land surface temperature for Adams county in Illinois captured on May 8th, 2011. Corn and soybean farmland images indicate cropland areas within the Adams county in Illinois in year 2011.

**Figure 4.**
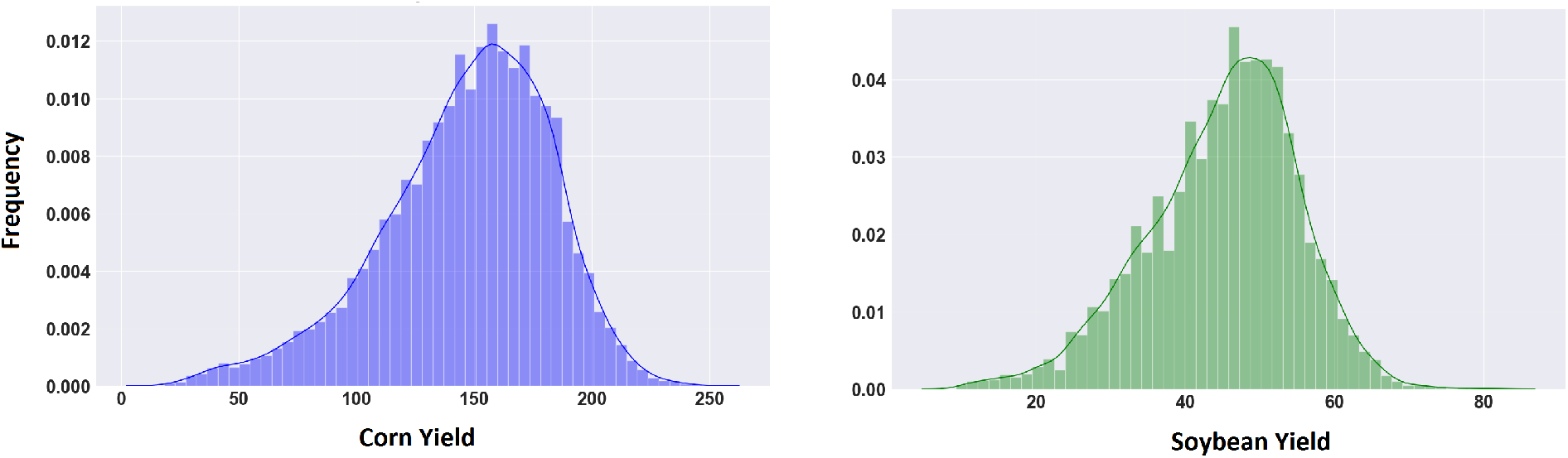
The histograms of the corn and soybean yields. The unit of yield is bushels per acre.

### 3.2 Design of Experiments

We compare our proposed method with the following models to evaluate the efficiency of our method:

#### Random forest (RF) [31]

RF is a non-parametric ensemble learning method which is robust against overfitting. We set the number and the maximum depth of trees in RF to be 150 and 20, respectively. We tried different number and the maximum depth of trees and found that these hyperparameters resulted in the most accurate predictions.

#### Deep feed forward neural network (DFNN)

DFNN is a highly nonlinear model which stacks multiple fully connected layers to learn the underlying functional relationship between inputs and outputs. The DFNN model has 9 hidden layers, each having 50 neurons. We used batch normalization for all layers. The ReLU activation function was used in all layers except the output layer. The model was trained for 120,000 iterations. Adam optimizer was used to minimize the Euclidean loss function.

#### Regression tree (RT) [32]

RT is a nonlinear model which does not make any assumption regarding the mapping function. We set the maximum depth of the regression tree to be 12 to decrease the overfitting.

#### Lasso [33]

Lasso adopts *L*_1_ norm to reduce the effect of unimportant features. Lasso model also can be used as a baseline model to compare the linear and nonlinear effect of input features. We set the *L*_1_ coefficient of Lasso model to be 0.05.

#### Ridge [34]

Ridge regression is similar to the Lasso model except it uses *L*_2_ norm as a penalty to reduce the effect of unimportant features. We set the *L*_2_ coefficient of Ridge model to be 0.05.

### 3.3 Training Details

YieldNet network was trained in an end-to-end manner. We initialized the network parameters with Xavier initialization [35]. To minimize the loss function defined in Equation (1), we used Adam optimizer [36] with learning rate of 0.05% and a mini-batch size of 32. The network was trained 4,000 iterations to convergence. We do not use dropout [37] because batch normalization also has a regularization effect on the network.

### 3.4 Final Results

After having trained all models, we evaluated the performances of our proposed method along with other competing models to predict corn and soybean yields. To completely evaluate all models, we took three years 2016, 2017, and 2018 as test years and predicted corn and soybean yields 4 times a year during the growing season on 23rd day of July, August, September, and October. Tables 3 and 4 present the corn and soybean yield prediction results, respectively, and compare the performances of models with respect to the root-mean-square error (RMSE) evaluation metric.

**Table 3:** The RMSE of corn yield prediction performance of models. The average ± standard deviation for corn yield in years 2016, 2017, and 2018 are, respectively, 165.72 ±30.35, 168.50 ± 32.88, and 170.77 ± 34.95. The numbers of test samples in years 2016, 2017, 2018 for corn yield prediction are 885, 882, and 784, respectively. The unit of RMSE is bushels per acre.

| Test Date |  | Models |  |  |  |  |  |
| --- | --- | --- | --- | --- | --- | --- | --- |
| Year | Month | Ridge | Lasso | RF | DFNN | RT | YieldNet |
| 2016 | July | 23.12 | 21.03 | 22.48 | 22.16 | 29.41 | **20.46** |
|  | August | 23.16 | 19.68 | 20.95 | 20.48 | 29.16 | **17.88** |
|  | September | 24.53 | 20.6 | 21.23 | 21.04 | 29.31 | **17.13** |
|  | October | 24.93 | 21.05 | 21.15 | 20.74 | 27.96 | **16.69** |
| 2017 | July | 30.55 | 27.53 | 26.61 | 26.40 | 33.64 | **22.73** |
|  | August | 25.16 | 22.27 | 22.25 | 20.85 | 28.02 | **18.44** |
|  | September | 24.15 | 21.5 | 21.99 | 19.21 | 26.8 | **16.11** |
|  | October | 25.73 | 20.94 | 22.14 | 18.90 | 26.78 | **16.74** |
| 2018 | July | 27.51 | 21.21 | 22.38 | 22.85 | 27.69 | **21.10** |
|  | August | 24.5 | 19.46 | 21.52 | 21.14 | 29.34 | **18.62** |
|  | September | 25.1 | 18.69 | 21.7 | 20.57 | 28.91 | **16.96** |
|  | October | 32.5 | 19.2 | 22.28 | 21.63 | 28.9 | **16.88** |
| Average |  | 25.91 | 21.10 | 22.22 | 21.33 | 28.83 | **18.31** |

**Table 4:** The RMSE of soybean yield prediction performance of models. The average ± standard deviation for corn yield in years 2016, 2017, and 2018 are, respectively, 53.94 ± 7.23, 50.24 ± 8.72, and 53.17 ± 9.72. The numbers of test samples in years 2016, 2017, and 2018 for soybean yield prediction are 773, 776, and 663, respectively. The unit of RMSE is bushels per acre.

| Test Date |  | Models |  |  |  |  |  |
| --- | --- | --- | --- | --- | --- | --- | --- |
| Year | Month | Ridge | Lasso | RF | DFNN | RT | YieldNet |
| 2016 | July | 9.87 | 9.12 | 9.38 | 8.27 | 10.82 | **5.32** |
|  | August | 8.2 | 8.42 | 9.15 | 6.89 | 10.65 | **5.25** |
|  | September | 8.3 | 8.27 | 9.12 | 7.19 | 10.37 | **4.90** |
|  | October | 8.49 | 8.85 | 9.02 | 7.62 | 10.7 | **4.44** |
| 2017 | July | 8.81 | 7.01 | 6.62 | 6.30 | 9.63 | **6.42** |
|  | August | 6.77 | 6.11 | 5.65 | 5.42 | 7.85 | **5.50** |
|  | September | 6.82 | 6.44 | 5.62 | 5.57 | 7.88 | **4.74** |
|  | October | 6.74 | 6.63 | 5.67 | 5.40 | 7.79 | **4.76** |
| 2018 | July | 8.77 | 9.88 | 8.13 | 8.62 | 9.74 | **7.04** |
|  | August | 7.75 | 8.08 | 7.78 | 7.14 | 9.58 | **6.36** |
|  | September | 8.25 | 8.04 | 7.78 | 7.53 | 9.64 | **5.22** |
|  | October | 8.01 | 8.14 | 7.9 | 7.41 | 9.4 | **5.50** |
| Average |  | 8.07 | 7.92 | 7.65 | 6.95 | 9.50 | **5.45** |

As shown in Tables 3 and 4, our proposed method outperforms other methods to varying extents. The Ridge and Lasso had comparable performances for soybean yield prediction, but, Lasso performed better compared to the Ridge for corn yield prediction. DFNN showed a better performance than RF, RT, and Ridge for both corn and soybean yield predictions. DFNN had a similar performance with Lasso for corn yield prediction while hiving higher prediction accuracy for soybean yield prediction. Despite the linear modeling structure, Lasso performed better than RF, and RT, which indicates that RF and RT cannot successfully capture the nonlinearity of remote sensing data, resulting in a poor performance compared to Lasso. RT had a weak performance compared to other methods due to being prone to overfitting. RF performs considerably better than RT because of using ensemble learning, which makes it robust against overfitting. Our proposed method significantly outperformed the other methods due to multiple factors: (1) the convolution operation in the YieldNet model captures both the temporal effect of remote sensing data collected over growing season and the spatial information of bins in histograms, (2) the YieldNet network uses transfer learning between corn and soybean yield predictions by sharing the weights of the backbone feature extractor, and (3) using a shared backbone feature extractor in the YieldNet model substantially decreases the number of model’s parameters and subsequently helps training process despite the limited labeled data.

The prediction accuracy decreases as we try to make prediction earlier during the growing season (e.g. July and August) due to loss of information. All models except our proposed model do not show a clear decreasing pattern in accuracy as we go from October to July, which indicates they cannot fully learn the functional mapping from satellite images to the yield.

We also report the yield prediction performance of our proposed model with respect to another evaluation metric, mean absolute error (MAE), in Table 5. As shown in Table 5, our proposed method accurately predicted corn yield one month, two months, three months, and four months before the harvest with MAE being 9.92%, 8.88%, 8.36%, and 7.8% of the average corn yield. The proposed model also accurately predicted soybean yield one month, two months, three months, and four months before harvest with MAE being 10.05%, 9.06%, 8.01%, and 7.67% of the average corn yield. The proposed model is slightly more accurate in soybean yield forecasting than corn yield forecasting, which is due to the higher variation in the corn yield compared to the soybean yield.

**Table 5:** The MAE of the corn and soybean yield prediction performances of our proposed model. The unit of MAE is bushels per acre.

| Crop | 2016 |  |  |  | 2017 |  |  |  | 2018 |  |  |  |
| --- | --- | --- | --- | --- | --- | --- | --- | --- | --- | --- | --- | --- |
|  | Jul. | Aug. | Sep. | Oct. | Jul. | Aug. | Sep. | Oct. | Jul. | Aug. | Sep. | Oct. |
| Corn | 14.92 | 14.32 | 14.36 | 14.48 | 18.24 | 15.92 | 12.71 | 13.40 | 16.11 | 14.59 | 13.35 | 13.23 |
| Soybean | 6.05 | 4.98 | 4.10 | 3.66 | 5.0 | 5.13 | 3.73 | 4.11 | 5.18 | 4.90 | 3.91 | 4.19 |

We visualized the error percentage maps for the corn and soybean yield predictions for the year 2018. As shown in Figures 5 and 6, the error percentage is below 5% for most counties, which indicates that our proposed model provides a robust and accurate yield prediction across US Corn Belt.

**Figure 5.**
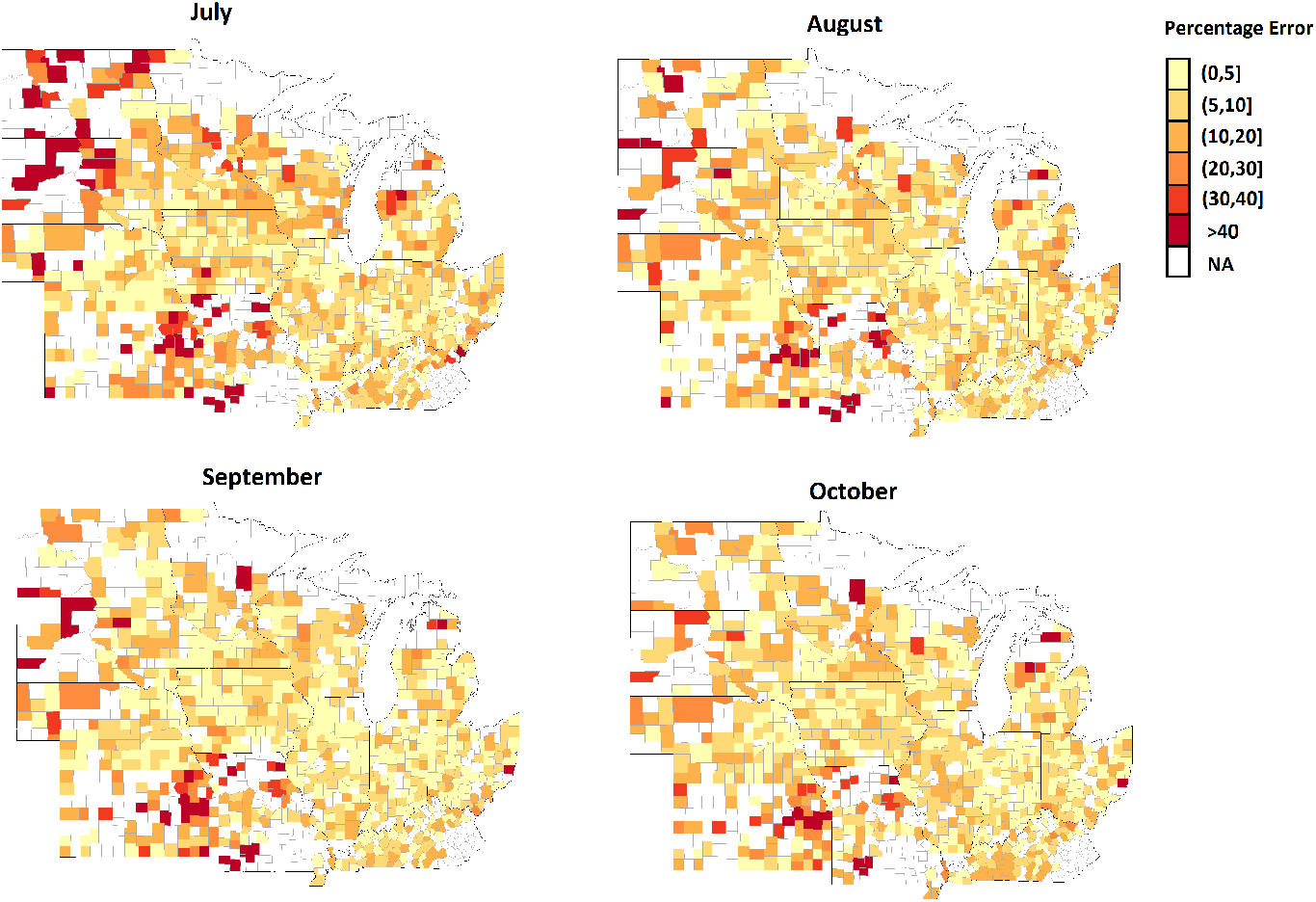
The error percentage maps for the 2018 corn yield prediction which is done during growing season in July, August, September, and October. The counties with white color indicate the ground truth yield were not available for those counties in 2018.

**Figure 6.**
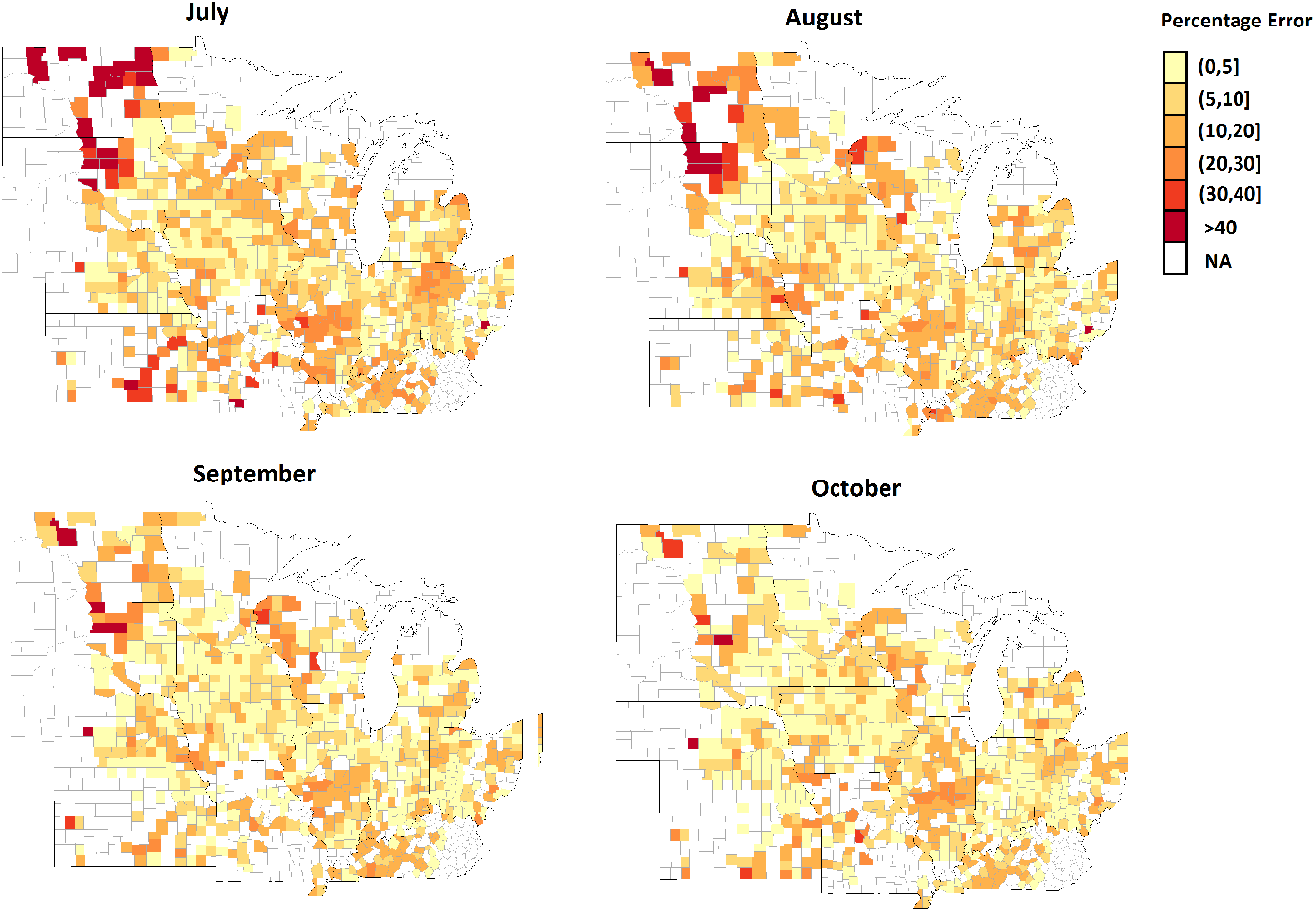
The error percentage maps for the 2018 soybean yield prediction which is done during growing season in July, August, September, and October. The counties with white color indicate the ground truth yield were not available for those counties in 2018.

To further evaluate the prediction results of our proposed model, we created the scatter plots of ground truth yield against predicted yield for the year 2018. Figure 7 depicts the scatter plots for the corn yield prediction during the growing season in months July, August, September, and October. The corn scatter plots indicate that the YieldNet model can successfully forecast yield months prior to the harvest.

**Figure 7.**
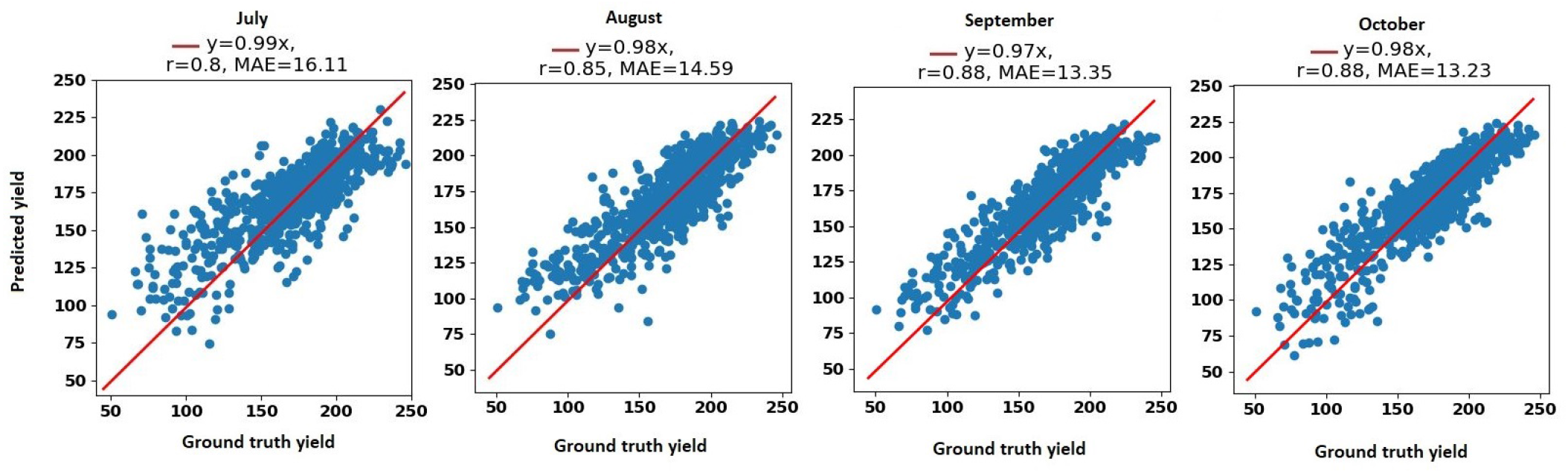
The scatter plots for the 2018 corn yield prediction during the growing season in months July, August, September, and October. MAE and r stand for mean absolute error and correlation coefficient, respectively. The unit of yield is bushels per acre.

Figure 8 depicts the scatter plots for the soybean yield prediction during the growing season in months July, August, September, and October. The corn scatter plots indicate that the YieldNet model provides reliable and accurate yield months prior to the harvest.

**Figure 8.**
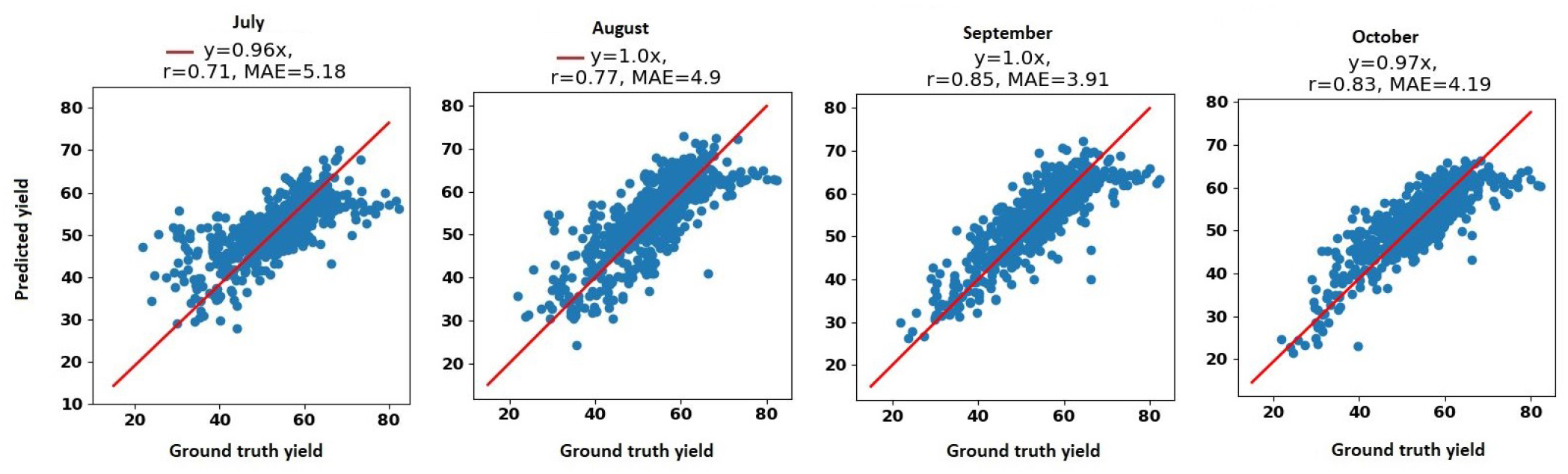
The scatter plots for the 2018 soybean yield prediction during the growing season in months July, August, September, and October. MAE and r stand for mean absolute error and correlation coefficient, respectively. The unit of yield is bushels per acre.

## 4 Ablation Study

In order to examine the usefulness of using a single deep learning model for simultaneously predicting the yield of two crops, we perform the following analysis. We train two separate models one for corn yield prediction and another for soybean yield prediction which are as follows:

### YieldNet^corn^

This model has exactly the same network architecture as the YieldNet model except we removed the the soybean head from the original YieldNet network. As a result, *YieldNet*^*corn*^ can only predict the corn yield.

### YieldNet^soy^

This model has exactly the same network architecture as the YieldNet model except we removed the the corn head from the original YieldNet network. As a result, *YieldNet*^*soy*^ can only predict the soybean yield.

Tables 6 and 7 compare the yield prediction performances of the above-mentioned models with the original YieldNet model.

**Table 6:**
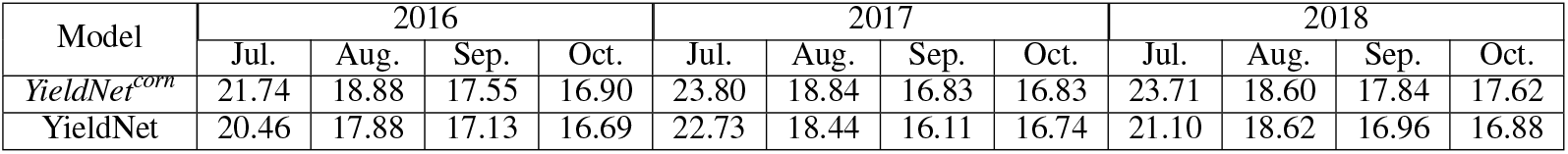
The RMSE of the corn yield prediction performances of the YieldNet and *YieldNet*^*corn*^ models. The unit of RMSE is bushels per acre.

**Table 7:** The RMSE of the soybean yield prediction performances of the YieldNet and *YieldNet*^*soy*^ models. The unit of yield is bushels per acre.

| Model | 2016 |  |  |  | 2017 |  |  |  | 2018 |  |  |  |
| --- | --- | --- | --- | --- | --- | --- | --- | --- | --- | --- | --- | --- |
|  | Jul. | Aug. | Sep. | Oct. | Jul. | Aug. | Sep. | Oct. | Jul. | Aug. | Sep. | Oct. |
| *YieldNet<sup>soy</sup>* | 6.49 | 5.49 | 5.15 | 4.57 | 6.96 | 5.82 | 4.86 | 4.86 | 7.80 | 6.58 | 5.46 | 5.57 |
| YieldNet | 5.32 | 5.25 | 4.90 | 4.44 | 6.42 | 5.50 | 4.74 | 4.76 | 7.04 | 6.36 | 5.22 | 5.50 |

As shown in Tables 6 and 7, the YieldNet model which simultaneously predicts corn and soybean yields outperforms individual *YieldNet*^*corn*^ and *YieldNet*^*soy*^ models. The YieldNet provides more robust and accurate yield predictions compared to the other two individual models, which indicates that transfer learning between corn and soybean yield prediction improves the yield prediction accuracy for both crops.

## 5 Discussion and Conclusion

Our numerical results illustrate that our approach to simultaneously predicting yield for both corn and soybeans is possible and can achieve higher accuracy than individual models. By utilizing transfer learning between corn and soybean yield to share the weights of the backbone feature extractor, YieldNet was able to substantially decrease the number of learning parameters. Our transfer learning approach enabled us to save on computation resources while also maximizing our prediction accuracy. Moreover, the accuracy achieve using a four month look-ahead has a lot of important implications for crop management decisions. With accurate yield predictions at various time points, decision makers now have the ability to change crop management practices to ensure yield is being maximized throughout its growth stage.

Although our approach highlighted corn and soybean in the U.S. market, this approach is applicable to any number of crops in any region. Due to the strength of our deep learning framework in combination with a generalized loss function, our approach is ready for scale. To improve the accuracy of our methodology, more data can be gathered and this can be left as a future extension to this work, along side more crops and more regions. It is the hope of this paper that our approach and results showcase the power deep learning for simultaneous yield prediction can have on the remote sensing community and the larger agricultural community as a whole.

